# Panton-Valentine leukocidin-induced neutrophil extracellular traps lack antimicrobial activity and are readily induced in patients with recurrent PVL^+^-*Staphylococcus aureus* infections

**DOI:** 10.1101/2021.11.26.470012

**Authors:** Hina Jhelum, Dora Čerina, CJ Harbort, Andreas Lindner, Leif Gunnar Hanitsch, Rasmus Leistner, Jennyver-Tabea Schröder, Horst von Bernuth, Miriam Songa Stegemann, Mariana Schürmann, Arturo Zychlinsky, Renate Krüger, Gerben Marsman

**Author notes:** Contributed equally to the manuscript (Renate Krüger, Gerben Marsman).

## Abstract

*Staphylococcus aureus* (*S. aureus*) strains that produce the toxin Panton-Valentine leukocidin (PVL; PVL-SA) frequently cause recurrent skin and soft tissue infections (SSTI). PVL binds to and kills human neutrophils, resulting in the formation of neutrophil extracellular traps, but the pathomechanism has not been extensively studied. Furthermore, it is unclear why some individuals colonized with PVL-SA suffer from recurring infections whereas others are asymptomatic. We thus aimed to (a) investigate how PVL exerts its pathogenicity on neutrophils and (b) identify factors that could help to explain the predisposition of patients with recurring infections.

We provide genetic and pharmacological evidence that PVL-induced NET formation is independent of NADPH-oxidase and reactive oxygen species (ROS) production. Moreover, through NET proteome analysis we identified that the protein content of PVL-induced NETs is different from NETs induced by mitogen or the microbial toxin nigericin. The abundance of the proteins cathelicidin (CAMP), elastase (NE), and proteinase 3 (PRTN3) was lower on PVL-induced NETs, and as such they were unable to kill *S. aureus.* Furthermore, we found that neutrophils from affected patients express higher levels of CD45, one of the PVL receptors, and are more susceptible to be killed at a low PVL concentration than control neutrophils. Neutrophils from patients that suffer from recurring PVL-positive infections may thus be more sensitive to PVL-induced NET formation, which might impair their ability to combat the infection.

**Importance:** Individuals colonized by *Staphylococcus aureus* strains that produce Panton-Valentine leukocidin (PVL-SA) often present with recurrent skin and soft-tissue infections, whilst other individuals remain asymptomatic. PVL is a toxin that kills neutrophils, which results in the formation of neutrophil extracellular traps. Traps induced by other stimuli are known to be toxic to *S. aureus.* We found however that NETs specifically induced by PVL are not toxic to *S. aureus*. Furthermore, we show that neutrophils from individuals that suffer from recurring PVL-SA infections are more sensitive to PVL-induced NET formation compared to healthy individuals. The significance of our work is in identifying a mechanism through which PVL-SA may actively counter the engagement of neutrophils. Moreover, we identified that patients with recuring PVL-SA infections may be more sensitive to this mechanism, which may help to explain their clinical condition and might provide avenues for future treatment development.

## Introduction

*S. aureus* is a widespread Gram-positive pathogen that causes infections in skin and soft tissue (SSTI), osteoarticular, airway, and the blood stream in humans and animals. Nasal colonization with *S. aureus* is found in up to 50% of humans, with strong age variation, and may be asymptomatic provided that skin and mucous membrane barriers are intact [1, 2].

Pathogenicity of *S. aureus* strains depends on a variety of virulence factors, among those, pore-forming leukocidins such as PVL that target neutrophils (also called polymorphonuclear leukocytes, PMN), and macrophages to evade phagocytosis and intracellular killing (reviewed in [3]). Over the last two decades an increasing number of outbreaks of SSTI in close communities [4–12] and in other health care settings [13,15,14,16–24] were caused by *S. aureus* strains that produce PVL (PVL-SA).

Although recurrent skin abscesses and furunculosis are often associated with PVL-SA infections, other pathologies are also linked to these strains. These include (a) worldwide reports of severe invasive infections - often complicated by thrombotic events - in previously healthy young individuals [25–27], (b) breast abscesses in lactating women [28] and transmission to offspring with life-threatening infections [29], and (c) necrotizing pneumonia [30], a severe manifestation, often fatal within days of hospital admission [31, 32]. In most cases, pneumonia is preceded by viral airway infections, like influenza, parainfluenza [31, 32] and SARS-CoV2 [33,35,34].

During outbreaks, identical PVL-SA strains have been obtained from individuals with asymptomatic nasal colonization as well as from patients with severe SSTI or invasive infections [11], suggesting that host factors may explain susceptibility to, and severity of, PVL-SA infections. Although studies in African Pygmies suggest that genetic variations in *C5ARI* may be associated with PVL-SA colonization [36], other factors that allow PVL-SA colonization or infections are not known.

The binding specificity of PVL defines both its host- and cell tropism. PVL is a two-component toxin consisting of LukF-PV and LukS-PV, which bind to the human panleukocyte receptor CD45 [37], and the human complement 5a receptors CD88/C5aR and C5L2, respectively [38]. As a result, human phagocytes are the major target of PVL. Upon binding, the subcomponents hetero-oligomerize into an octameric membrane spanning pore [39, 40], which drives cell death. Neutrophils express higher levels of both C5a receptors than monocytes [38] and are more sensitive to PVL mediated killing than both monocytes and macrophages [41], suggesting they are the major target of PVL. Importantly, neutrophils are essential for host defense against *S. aureus* [42] and patients with impaired production of reactive oxygen species (ROS) in phagocytes are particularly sensitive to *S. aureus* infections (see review [43]). Neutrophils clear invading *S. aureus* through phagocytosis and by undergoing NETosis, a cell death process which results in the formation of neutrophil extracellular traps (NETs). NETs consist of an externalized web of chromatin decorated with antimicrobial peptides and proteases and are toxic to bacteria [44].

Depending on the stimulus, NET formation may require ROS formation [45]. A common mediator of ROS-dependent NET formation is NADPH-oxidase, which produces superoxide (O^2-^) that spontaneously dismutates to hydrogen peroxide (H2O2). Myeloperoxidase (MPO) converts hydrogen peroxide into highly reactive hypochlorous acid (HOCl). ROS production allows for the release of neutrophil proteases from granules, protease-mediated chromatin decondensation and the final lysis of the plasma membrane [46–48]. Two *S. aureus* toxins that kill neutrophils and result in NET formation are γ-hemolysin AB and PVL [49–51]. Interestingly, *S. aureus* also produces nucleases to escape from NETs [52]. Whether NET formation is a beneficial or detrimental neutrophil response, both for the host as well as for *S. aureus*, remains unclear and may be context dependent [50].

In this study, we set out to characterize PVL-induced NET formation with the aim to understand its possible contribution to the pathogenesis of PVL-SA infections. We show that PVL-induced NET formation is NADPH-oxidase independent. Consistently, neutrophils isolated from chronic granulomatous disease (CGD) patients, which have a mutation in the gene encoding one subunit of the NADPH oxidase complex, make NETs in response to this toxin [53]. PVL-induced NETs showed quantitative proteomic differences to NETs produced after PMA or nigericin stimulation. PVL-NETs contained a lower abundance of multiple antimicrobial proteins and in contrast to PMA- or nigericin-NETs, did not kill *S. aureus.* Given the lack of antimicrobial activity, we asked whether neutrophils from patients with a history of recurrent PVL-SA infection show altered sensitivity to PVL compared to controls. Indeed, we found that neutrophils from these patients express higher levels of the PVL receptor CD45 than healthy controls and make more NETs in response to a low concentration of PVL. Our results suggest that PVL-receptor expression may mediate susceptibility to symptomatic *S. aureus* infection and that NET-formation induced by this toxin serves as an offensive strategy to preemptively neutralize the neutrophil.

## Results

### PVL induces NETs independent from ROS

To further our understanding of the role of PVL toxin in PVL-SA pathogenesis, we verified [50, 51] and further characterized PVL-induced NET formation. NADPH-oxidase dependent superoxide formation is essential in NETs induced by some stimuli [45], prompting us to investigate the NETosis pathway initiated by PVL. We first tested superoxide generation in healthy primary neutrophils stimulated with different concentrations of PVL or PMA, a well characterized mitogen that induces NADPH-oxidase dependent NETosis [54]. PVL stimulation resulted in weak superoxide production at all concentrations (Fig 1A). In contrast, PMA induced a significant superoxide burst.

**Fig 1.**
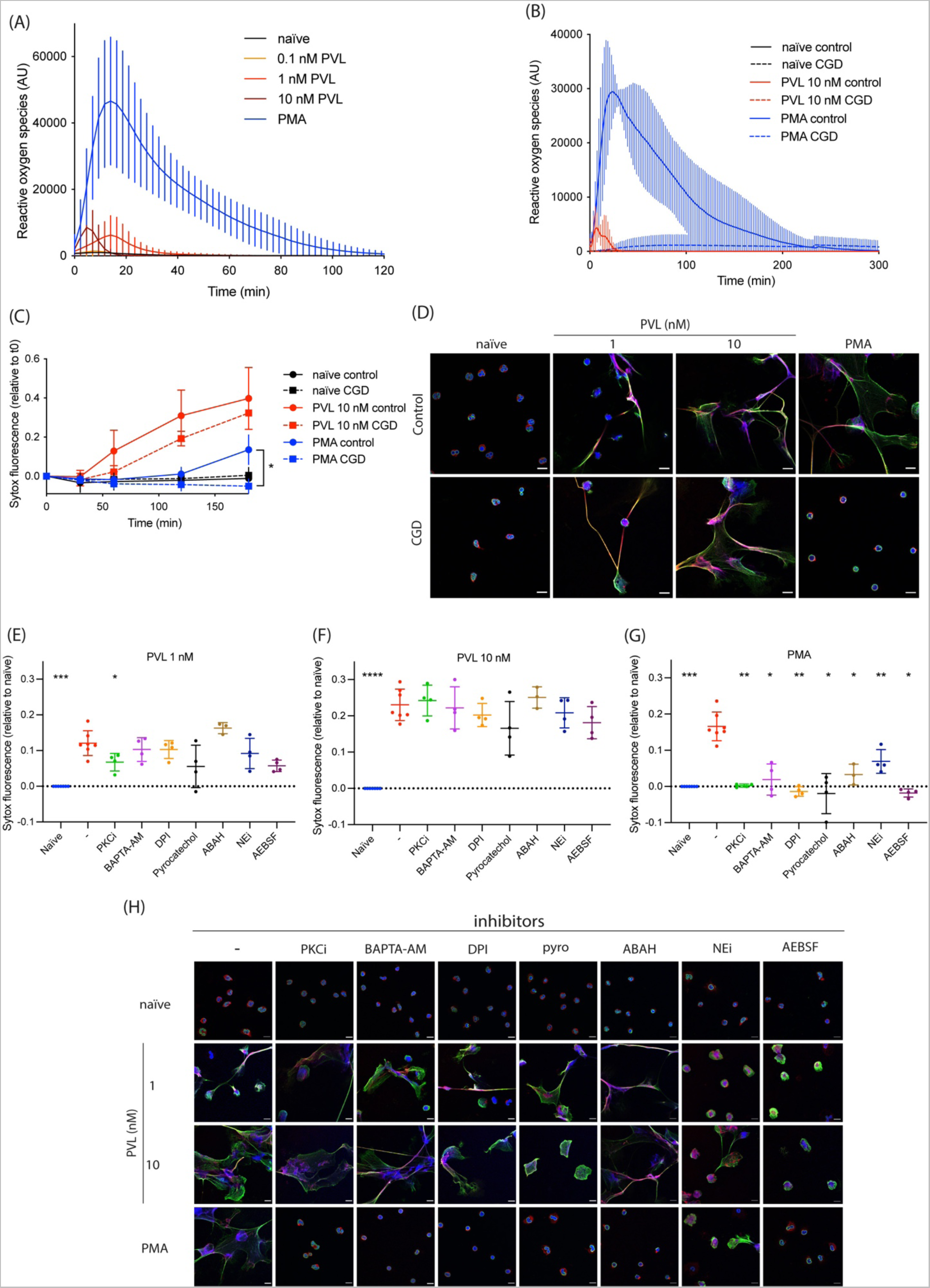
PVL induces ROS-independent NET formation. (A-B) Reactive oxygen species (ROS) formation was quantified in control (A n=14, B n=3) or CGD human primary neutrophils (n=3) stimulated with PMA (50 nM) or PVL (0.1 nM, 1 nM or 10 nM). (C) Control or CGD neutrophils (n=3) were stimulated with PMA (50 nM) or PVL (10 nM) for 3h and cell death was quantified using the cell impermeable DNA dye SYTOX Green. Indicated is SYTOX Green fluorescence relative to *t* = 0 min. (D) Representative confocal microscopy images of control and CGD patient neutrophils either naïve or stimulated with PMA (50 nM) or PVL (1 nM or 10 nM) and stained for DNA (blue), NE (red) and chromatin (green). Scale bar represents 10 µm. (E-G) Healthy human neutrophils were treated with PKC inhibitor Gö6983 (PKCi, 1 μM), BAPTA-AM (10 μM), DPI (1 μM), pyrocatechol (30 μM), ABAH (500 μM), NEi (20 μM) and AEBSF (100 μM) for 30 min, before stimulating with (E) PVL 1 nM, (F) PVL 10 nM or (G) PMA 50 nM for 4h. We quantified cell death with SYTOX Green and the fluorescence signal relative to naïve is indicated. (H) Representative confocal microscopy of naïve neutrophils or after stimulation with PMA or PVL (1 nM or 10 nM) in the presence or absence of indicated inhibitors, and stained for DNA (blue), NE (red) and chromatin (green). Scale bar represents 10 µm. (C) Two-way ANOVA with Tukey’s post-hoc test. \**P*<0.05. (E-G) Mean ±SD of four independent experiments. \**P*<0.05, ** *P*<0.01, *** *P*<0.001, one-way ANOVA with Dunnett’s multiple comparison test.

We quantified ROS production as well as NET formation in neutrophils from CGD patients. As expected, CGD neutrophils failed to generate superoxide in response to PVL or PMA (Fig 1B). However, unlike PMA, PVL induced extracellular DNA release (Fig 1C) and NET formation (Fig 1D and S1-S8 movies) in CGD neutrophils. These data show that while PVL stimulation weakly activates NADPH-oxidase, NET formation is independent from the produced ROS.

Given the known involvement of various neutrophil proteins in NET formation, we asked if PVL-induced NET formation could be blocked pharmacologically by the PKC inhibitor Gö6983 (PKCi), the calcium chelator BAPTA-AM, the NADPH-oxidase inhibitor DPI, the ROS scavenger pyrocatechol, the myeloperoxidase inhibitor ABAH, the neutrophil elastase inhibitor NEi, or the pan serine protease inhibitor AEBSF (Fig 1E-F and 1H). We observed that PKCi weakly inhibited NET formation induced by 1 nM PVL (Fig 1E), whilst none of the other inhibitors, including inhibitors of ROS-producing enzymes (DPI and ABAH) or ROS scavengers (pyrocatechol), blocked NET formation induced by 1 or 10 nM PVL (Fig 1E-F and 1H). Notably, neutrophils pre-treated with NEi or AEBSF and stimulated with 1nM PVL formed smaller NETs (Fig 1H). These data suggest that PVL-mediated NET formation is ROS- and protease-independent. As a control, we confirmed [45] that PMA-induced NET formation was blocked by all inhibitors tested (Fig 1G-H).

Mazzoleni *et al.* [51] suggested that small conductance calcium-activated potassium (SK) channels and alternative ROS sources, such as mitochondria or xanthine oxidase, mediate PVL-induced NET formation. To verify this, we pre-treated neutrophils with the xanthine oxidase inhibitor allopurinol, the SK channel inhibitor apamin, and the mitochondrial uncouplers DNP and FCCP, before stimulating with either PMA or PVL. The SK channel inhibitor NS8593 was toxic to neutrophils in our hands and was excluded from our experiments. At 1 nM, but not at 10 nM PVL stimulation, allopurinol inhibited NET formation, whilst mitochondrial uncoupling with FCCP partially inhibited NET formation. (S1A-B Fig). These findings were supported by confocal imaging (S1D Fig). None of the inhibitors affected PMA-induced NET formation (S1C Fig). Given that PVL induces NET formation in CGD patients despite a complete lack of ROS (Fig 1B), and the inability of the ROS scavenger pyrocatechol to inhibit PVL-induced NET formation, the inhibition of NETs by allopurinol treatment or mitochondrial uncoupling appears to be independent of ROS production. This is further supported by the observation that DPI, which not only inhibits NADPH-oxidase but all flavin-containing proteins including xanthine oxidase [55] and mitochondrial complex I [56], did not block PVL-induced NET formation (Fig 1E-F). Finally, the inability of ROS-targeting inhibitors to block NET formation induced by 10 nM PVL indicates that PVL-induced NET formation is independent from ROS production.

### PVL-induced NETs lack enrichment of key antimicrobial proteins

Since PVL induces NETs through a non-canonical pathway we hypothesized that the resulting NETs may have different protein compositions. We therefore analyzed the proteomes of NETs induced by PVL, PMA, and the NADPH-oxidase independent NET inducer nigericin [45]. We analyzed NETs from three independent donors by quantitative mass spectrometry and compared them to the proteome of unstimulated neutrophils. Principal component analysis revealed that PVL-, PMA- and nigericin-induced NETs each have distinct proteomes (Fig 2A). Interestingly, PVL-NETs appear to cluster between naïve neutrophils and NETs induced by PMA and nigericin, suggesting their proteome composition is intermediate between naïve cells and classical NETs. Measurement of the Euclidean distance between samples confirmed that the PVL-NET proteome is less distinct from naïve neutrophils than that of PMA- or nigericin-induced NETs (S2A Fig).

**Fig 2.**
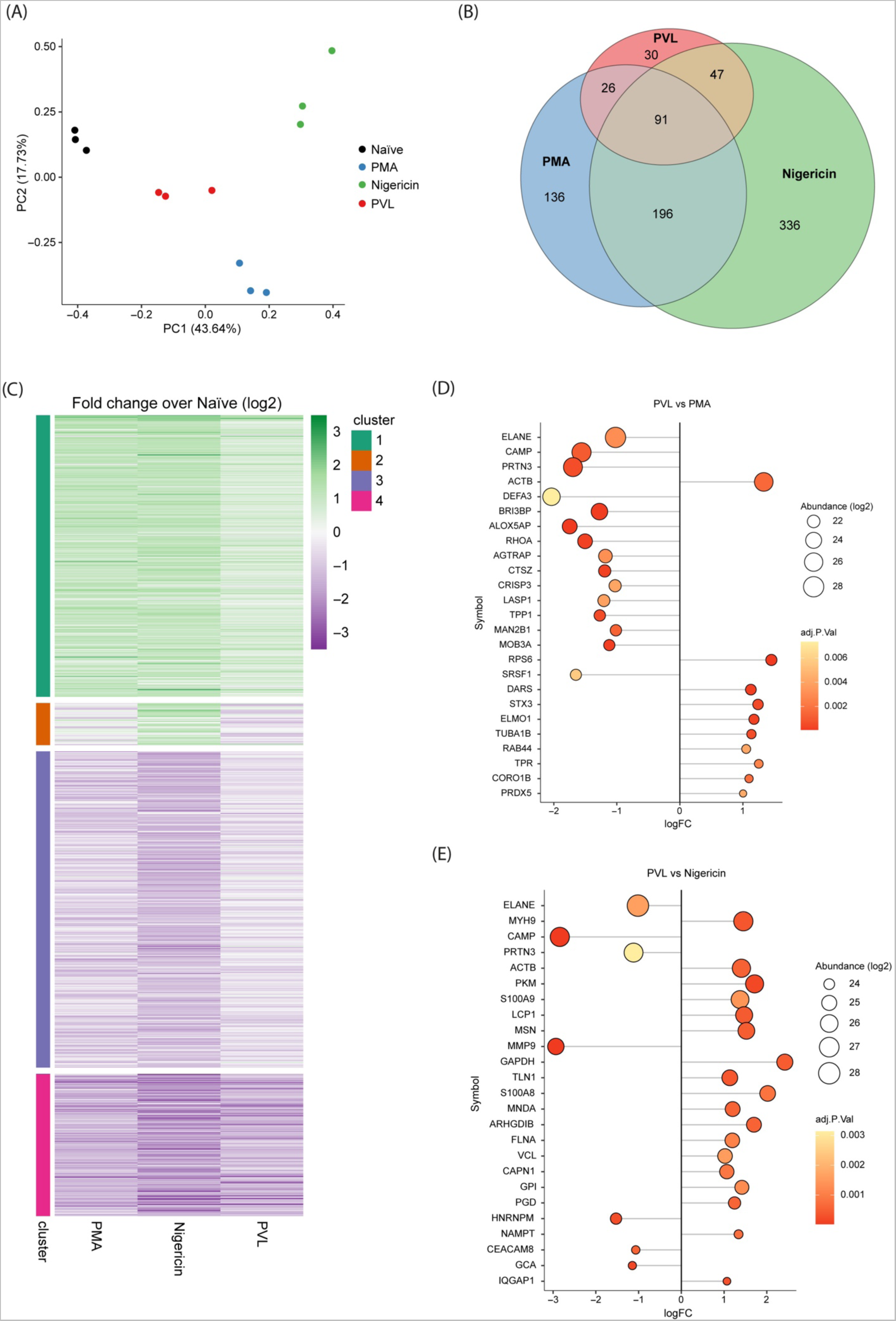
NET proteome assembly is different in PVL-induced NETs. (A) Principal component analysis of the proteomes of naïve neutrophils, and PVL-, PMA-, and nigericin-induced NETs. Analysis was performed on scaled log2-abundance of all proteins detected in at least 2/3 replicates. (B) Euler diagram showing the number and distribution of significant Differentially Abundant Peptides (logFC > 1, adjusted *p* < 0.01) on NETs compared to naïve neutrophils. Areas are proportional to DAP set size. (C) Clustered heatmap showing fold change (log2) values on each NET sample compared to naïve neutrophils of significant DAPs from (B). Clustering was performed by k-means algorithm with k=4 clusters. (D-E) Top 25 most-abundant DAPs that are significantly differentially abundant on PVL-induced NETs compared with PMA-(D) and nigericin-induced NETs (E). Point size and fill color represent average abundance across samples and adjusted *p* value, respectively. Proteomes were made from n=3 samples per condition from independent donors. DAP significance for each comparison was determined by a threshold of |[log2 fold change]| > 1 and adjusted *p* value < 0.01.

We detected 2458 distinct proteins in the combined NET samples (present in at least two of three replicate experiments). To examine which proteins are driving the differences between the NET proteomes, we performed a differential enrichment analysis using a 2-fold change in protein abundance with an adjusted *p* value below 0.01 as a cut-off. We found significant differences in relative protein abundance between all three NET stimuli when compared with naïve neutrophils (Fig 2B and S2A Fig). Fewer differentially abundant proteins (DAPs) were detected between PVL-induced NETs and naïve PMNs (194 significant) compared to PMA- (449) and nigericin-induced NETs (670). These data suggest that the specific enrichment or depletion of proteins during NET formation is lower or incomplete in PVL-treated neutrophils.

Examination of the fold changes of NET-specific DAPs revealed that most DAPs have the same pattern of enrichment or depletion across all types of NETs compared to naïve neutrophils (459 with negative logFC, 294 positive logFC, Fig 2C). Regression of global DAP fold changes (Fig S2B) revealed that though most proteins have the same enrichment pattern across NETs, the fold-changes are reduced in PVL-compared to PMA- and nigericin-induced NETs (slopes = 0.64, and 0.49 respectively), while PMA and nigericin NETs have equivalent DAP fold-changes (slope = 1).

K-means clustering of DAP fold change patterns supported this trend, and further identified a cluster of proteins specifically enriched on nigericin induced NETs (Fig 2C cluster two, orange, and S1 Table). To look more closely at the differences between PVL-induced NETs and PMA- or nigericin-induced NETs, we performed pair-wise comparisons between PVL abundant DAPs (Fig 2D and 2E, respectively). Among these, key neutrophil proteins such a NE (ELANE), PRTN3, and CAMP, were all less abundant on PVL-induced NETs compared with PMA- or nigericin-induced NETs. In contrast, we found cytoskeleton proteins such as actin (ACTB), myosin (MYH9), and tubulin (TUBA1B) to be more enriched on PVL-NETs. Our data suggest that PVL induces NETs with a different NET proteome assembly when compared with PMA or nigericin, resulting in a lack of enrichment of key antimicrobial proteins on PVL-NETs.

### PVL-induced NETs are less bactericidal

Given the differential enrichment of antimicrobial proteins in PVL-compared to PMA- and nigericin-NETs, we hypothesized that their bactericidal activity may be different. We incubated methicillin-resistant *S. aureus* (MRSA) with either PVL-, PMA-, or nigericin-induced NETs for 1 h, harvested the bacteria, and quantified the surviving colony forming units. PVL-induced NETs did not kill MRSA, whilst PMA- and nigericin-induced NETs did (Fig 3A). We verified that all stimuli at these concentrations released similar amounts of NETs by nanodrop DNA quantification (Fig 3B). To exclude a possible contribution of killing through phagocytosis we performed the same experiment in the presence of cytochalasin B. We did not observe a difference in antimicrobial activity upon inhibition of phagocytosis, indicating that NETs were solely responsible for the killing (Fig S3). These results show that PVL-NETs have a lower antimicrobial potential against MRSA than PMA- or nigericin-NETs.

**Fig 3.**
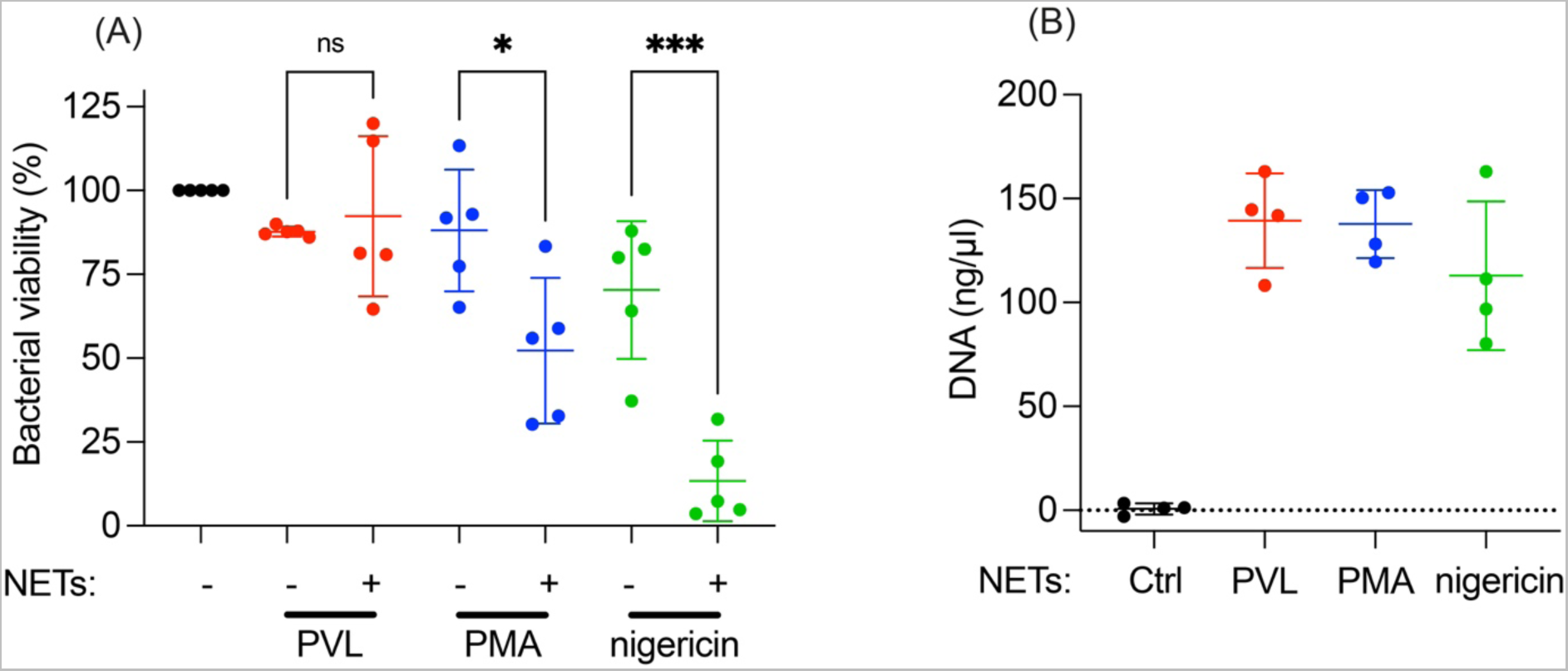
PVL-induced NETs do not kill MRSA. (A) Primary human neutrophils were stimulated with PVL (100 nM), PMA (100 nM), or nigericin (30 µM), for 4 h to induce NETs. MRSA was incubated with NETs for 1 h and CFU was quantified. Bacterial viability is expressed as a percentage of CFUs normalized to MRSA incubated in the absence of NETs. (B) The DNA content of PVL-, PMA-, and nigericin-induced NETs at 4 h was quantified by spectrophotometry. (A) Mean ± SD of five independent experiments. *p<0.05, *** p<0.001, one-way ANOVA. (B) Mean ± SD of four independent experiments.

### PVL induces NETs efficiently in neutrophils from patients with recurrent PVL-SA infections

The lack of antimicrobial activity of PVL-NETs prompted us to ask whether neutrophils from patients that suffer from recurrent PVL-SA infections show altered responses to PVL that may help to explain the patients’ increased susceptibility to infection. We quantified NET formation in response to 0.1 nM, 1 nM, and 10 nM PVL in patients and controls using a cell impermeable DNA dye. Interestingly, 0.1 nM PVL killed neutrophils isolated from patients more efficiently than neutrophils isolated from healthy controls (Fig 4A). Patient and control neutrophils were equally susceptible to make NETs in response to PMA.

**Fig 4.**
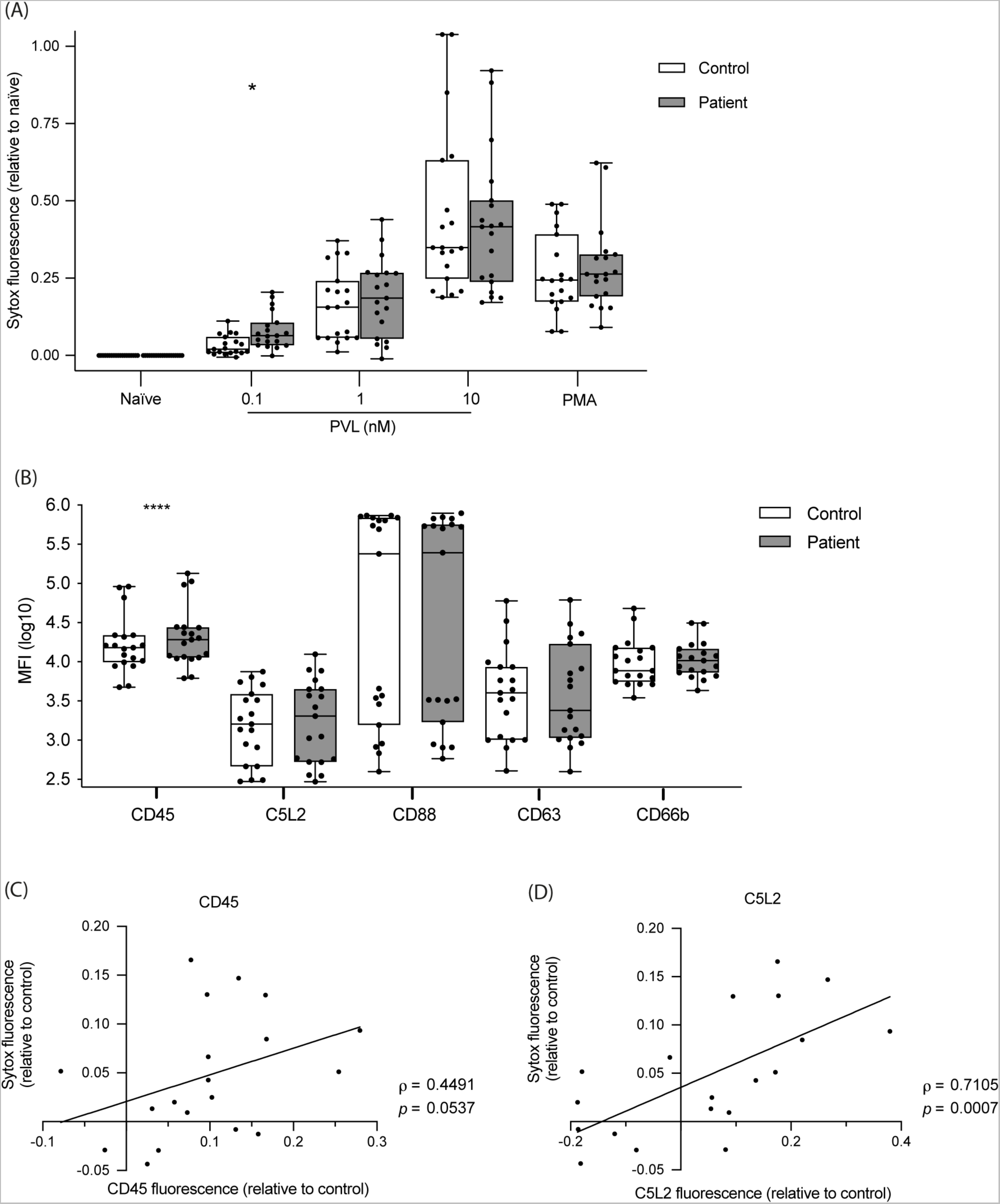
Neutrophils from patients with recurrent PVL-SA infections make more NETs in response to PVL and express more CD45 than neutrophils from healthy donors. (A) Purified primary neutrophils from patients suffering from recurrent PVL-SA infections (n=19) and healthy controls (n=19) were either left untreated or stimulated with PMA (50 nM) or PVL (0.1 nM, 1 nM or 10 nM) for 4 h and cell death was quantified by adding the cell-impermeable DNA dye SYTOX Green and measuring fluorescence. Boxplot indicates SYTOX fluorescence relative to naïve neutrophils at 4 h. (B) CD45, C5L2, CD88, CD63 and CD66b were immunolabelled on neutrophils from patients and healthy controls. The fluorescence was measured and indicated as log-transformed MFI. (C) CD45 or (D) C5L2 fluorescence of patient neutrophils relative to control was plotted against SYTOX fluorescence of patient neutrophils relative to control after incubation with 0.1 nM PVL for 4 h. (A) Unpaired Wilcoxon signed rank test with Bonferroni post-hoc test, * *P*<0.05. (B) Paired *t*-test, *** *P*<0.001. (C-D) Spearman’s correlation test where the coefficient of correlation (ρ) and probability (p) are indicated. A best-fit line indicates the trend.

Given the binding specificity of PVL we checked the expression of CD45, CD88, and C5L2, as well as the neutrophil activation markers CD63 and CD66b on control and patient neutrophils by flow cytometry. Patient neutrophils expressed higher levels of CD45 (adjusted *p* value = 0.000473), and there was no difference in CD88. Furthermore, a subset of patients expressed increased C5L2 levels compared to control neutrophils (Fig S4A), but this trend was not consistent for all patients (Fig 4B). Interestingly, the difference in expression of C5L2 and possibly CD45 in patients compared to controls correlated with the difference in NET formation in response to 0.1 nM PVL (ρ = 0.4491 for CD45, ρ = 0.7105 for C5L2) (Fig 4C and 4D). Differences in expression levels of CD88, CD63 or CD66b did not correlate with differences in NET formation (Fig S4B-D).

Taken together, our data suggest that neutrophils from patients with recurrent PVL-SA infections express higher levels of the PVL receptor CD45 and may be more sensitive to PVL-induced NET formation at low PVL concentrations when compared to control neutrophils.

## Discussion

In the past few decades, PVL has emerged as an important virulence factor in community acquired SA infections. Interestingly, human host factors that mediate susceptibility or severity of PVL-SA infections are not known. To further our understanding of the contribution of PVL to PVL-SA pathogenesis, we set out to characterize PVL-induced NET formation. PMA or *Candida albicans* induce NET formation which is ROS-dependent and is blocked by compounds that scavenge ROS or inhibit ROS [45]. Other stimuli, like nigericin and calcium ionophores induce NETs through a ROS independent mechanism [45]. We identified that PVL induces the formation of NETs independently of NADPH-oxidase and MPO activity, two enzymes that produce ROS. This is evidenced by two observations (1) CGD neutrophils do not produce ROS upon stimulation with PVL, but make NETs, and (2) DPI, an inhibitor of flavin-containing proteins such as NADPH-oxidase, xanthine oxidase [55], and cytochrome C [57, 56], the ROS scavenger pyrocatechol, and the MPO inhibitor ABAH all fail to inhibit PVL-induced NET formation.

A recent study by Mazzoleni *et al*. suggested that ROS derived from mitochondria or xanthine oxidase are involved in PVL-induced NET formation [51]. We observed that treatment with DNP or FCCP, as well as xanthine oxidase inhibition by allopurinol partly inhibited PVL-induced NET formation. Xanthine oxidoreductase consists of two isoforms, xanthine oxidase and xanthine dehydrogenase. Allopurinol inhibits both isoforms, which affects purine metabolism. Moreover, allopurinol itself generates superoxide radicals upon inhibition of xanthine oxidase [58]. Therefore, its inhibitory effect on PVL-induced NET formation may be independent from xanthine oxidase inhibition or ROS formation. How allopurinol exerts its inhibitory effects on PVL-induced NET formation is still unclear.

Furthermore, given that both DPI and the ROS scavenger pyrocatechol were unable to inhibit PVL-induced NET formation, we hypothesize that the inhibitory effect of allopurinol and FCCP is independent from their effects on ROS production. Notably, inhibition of NET formation by these inhibitors was seen at 1 nM PVL but was lost when stimulating with 10 nM PVL. It may be that PVL at high concentrations is able to induce NET formation without the involvement of neutrophil components interrogated by the inhibitors used in our study, whilst at low PVL concentrations some components may contribute to NET formation.

Neutrophil proteases were not essential for PVL-induced NET formation, although we did observe that formed NETs appeared smaller upon protease inhibition. Neutrophil elastase is known to contribute to chromatin decondensation during NADPH-oxidase dependent NETosis [47, 59] and this decondensation may occur during PVL-mediated NET formation as well. Although PAD4-mediated histone citrullination has been suggested to be involved in NADPH oxidase-independent NET formation, others have excluded its involvement in driving PVL-induced NET formation [51].

Interestingly, different NETosis mechanisms lead to NETs with specific compositions. We find differences in protein abundances of NETs induced by PVL, PMA, or nigericin, and the proteome of PVL NETs specifically appears to be less distinct from that of unstimulated neutrophils. Several antimicrobial proteins are less abundant in PVL-induced NETs, when compared to PMA- or nigericin-induced NETs. In contrast, cytoskeletal proteins are more abundant on PVL NETs. A previous report also identified a similar enrichment of cytoskeletal proteins in spontaneously lysed neutrophils [60]. Given PMA or nigericin-induced NET formation involves an intracellular release of granular components [59], this ‘mixing-of-the- bag’ may allow for efficient tethering of granular components to the chromatin backbone and degradation of the cytoskeleton. Cell lysis occurs 2-3 h after stimulation with PMA, while PVL-induced NET formation occurs swiftly, with cell lysis occurring within the first 2 h after stimulation. We speculate that granular components may be lost to the extracellular environment during this rapid lysis, resulting in a less bactericidal NET. These data suggest that NETs induced by different stimuli may display different functional characteristics. Leukocidins such as PVL may have evolved to elicit a harmless form of NET formation to promote survival of the invading bacteria.

We hypothesized that the lack of antimicrobial activity of PVL-NETs might contribute to PVL-SA pathogenesis and asked whether neutrophils from patients that suffer from recurrent PVL-SA infections show unusual responses to PVL. We identified increased NET formation in patient neutrophils upon stimulation with a low PVL concentration, and a higher expression of CD45 and potentially C5L2, two of the toxin’s receptors, in patient neutrophils compared to healthy controls. Furthermore, the increase in CD45 and C5L2 expression correlated with more NET formation induced by PVL.

LukS-PV and LukF-PV have different binding affinities to their respective receptors (C5L2 and CD88 for LukS-PV, CD45 for LukF-PV), and the low binding affinity of LukF-PV to CD45 suggests that varying expression levels of CD45 are unlikely to modulate pore formation [37]. In contrast, LukS-PV binding affinity to CD88 or C5L2 is much higher and therefore their expression may modulate sensitivity to PVL toxin. Given that neutrophils express C5L2 less abundantly than CD88, it was surprising to find that differential expression of C5L2 correlates with an increased sensitivity of patient neutrophils to PVL [38]. However, the affinities of LukS-PV binding to CD88 and C5L2 have never been compared and the relative contribution of C5L2-binding to PVL mediated cell death is therefore not known. Whether there is a genetic predisposition in patients with recurrent PVL-SA remains to be determined and we are currently investigating avenues to study this.

CD88 localizes to the plasma membrane, rendering it accessible for extracellular PVL toxin. However, there are conflicting reports on the localization of C5L2 since it has been detected both intracellularly and on the plasma membrane [61, 62]. We detected C5L2 on the plasma membrane through flow cytometry, suggesting that the expression is not exclusively intracellular. However, much is unknown concerning the regulation of C5L2 and CD88 expression and receptor shuttling and it is unclear how this regulation might affect targeting by PVL [61, 62].

Pore formation by PVL is described to exert various cellular effects such as intracellular calcium flux, ATP-release into the extracellular environment, induction of apoptosis and cellular lysis [63]. These various phenotypes likely depend on the toxin concentration a cell encounters. We therefore characterized NET formation at different PVL concentrations and observed robust NET formation upwards of 1 nM PVL. In clinical samples from PVL-SA patients, median levels of PVL toxin were previously found to be 0.42µg/ml (∼11 nM) with a range of 0-399 µg/ml, which suggests that NET induction may be expected in these patients [64], notwithstanding the difficulty of predicting PVL potency *in vivo*.

NETs have at least three described functions, they are antimicrobial [44], procoagulant [65], and activate the immune system [66, 67]. PVL-SA infections may be associated with a higher risk for thrombotic events. In addition to being less antimicrobial, it would be interesting to investigate if PVL-induced NETs also differ in their procoagulant or immune activation functions.

In conclusion, our observation of (a) differentially expressed PVL receptors in individuals with recurrent or severe PVL-SA infections and (b) functionally different NET formation after neutrophil exposure to PVL in contrast to other stimuli may explain specific clinical features and interindividual differences in PVL-SA infections. We propose that overexpression of PVL receptors might make an individual’s neutrophils more prone to this disarmed NET formation, preventing efficient clearance of the invading bacteria. In turn, this could provide a competitive advantage to PVL-SA, facilitating colonization and recurrent infections. Further investigations are necessary to verify this finding and find potential treatment or prevention strategies for these infections.

## Materials and Methods

### Neutrophil isolation, experimental conditions, and inhibitors

Human neutrophils were isolated by layering whole blood over Histopaque-1119 (Sigma) followed by a discontinuous Percoll gradient (Amersham Biosciences) as previously described [46]. Alternatively, they were also isolated using the direct human neutrophil isolation kit (EasySep, StemCell Technologies) following manufactureŕs instructions.

All experiments were performed in Seahorse XF RPMI medium (SF-RPMI, Agilent) supplemented with 2 mM glutamine, 10 mM HEPES, 1 mM glucose and 0.1% human serum albumin (HSA) at pH 7.4 except where mentioned. The stimuli used to induce NET formation were phorbol 12-myristate 13-acetate (PMA, Sigma), PVL (equal amounts of *S. aureus* recombinant LukS and LukF (Bioservices), and nigericin (InvivoGen). We used the following inhibitors: Gö6983 (PKCi, Tocris) BAPTA-AM (Thermo Fisher Scientific), pyrocatechol (Sigma), 4-Aminobenzoic acid hydrazide (ABAH, Cayman chemical), neutrophil elastase inhibitor (NEi, MedChemExpress), Diphenyleneiodonium chloride (DPI, Calbiochem), AEBSF (Sigma), allopurinol (Sigma), Apamin (Sigma), 2,4-Dinitrophenol (DNP, Sigma) and 2-[[4-(trifluoromathoxy) phenyl]hydrazinylidene] propanedinitrile (FCCP, Abcam).

### ROS measurement

Purified neutrophils were seeded at 10^5^ cells per well in a 96-well plate and incubated for 30 min with the indicated inhibitors, followed by incubation for 10 min with 50 μM luminol and 1.2 units/ml horseradish peroxidase at 37°C prior to stimulation with indicated stimuli. Luminescence was measured over time in a VICTORX luminometer (Perkin Elmer) [45].

### Neutrophil lytic cell death assay

10^5^ neutrophils were seeded in a 96-well plate and incubated with the appropriate inhibitors for 30 min, followed by incubation with 50 nM cell-impermeable DNA dye SYTOX Green (ThermoFisher Scientific) for 5 min at 37°C, prior to stimulation with indicated stimuli. Fluorescence was recorded once per hour for 4h using a Fluoroskan Ascent (ThermoFisher Scientific).

### NET staining

10^5^ neutrophils were seeded on glass coverslips in a 24-well plate and incubated with or without appropriate inhibitors followed by stimulation with indicated stimuli for 4 h at 37°C. Cells were fixed with 4% paraformaldehyde (PFA) for 20 min at room temperature. Cells were washed with PBS, permeabilized with 0.5% Triton X-100 for 1 min and incubated in blocking buffer (3% normal goat serum, 3% cold water fish gelatin, 1% bovine serum albumin and 0.05% Tween-20 in PBS) for 30 min. Neutrophils were then stained using antibodies detecting elastase (Merck Millipore, 481001) and a subnucleosomal complex of Histone 2A, histone 2B and chromatin [68]. The secondary antibodies used were goat anti-mouse Alexa Fluor 488 (Invitrogen, A11029) and goat anti-rabbit Alexa Fluor 647 (Invitrogen, A21245) followed by staining with DAPI (0.1 μg/ml, Invitrogen). Finally, the samples were mounted using antifade mountant (ProLong Diamond Antifade mountant, ThermoFisher Scientific). Images were acquired using a Leica TCS SP8 confocal microscope.

For live NET imaging, cells were resuspended in Agilent XF RPMI medium (Agilent) at pH 7.4, supplemented with 0.1% Human Serum Albumin, 10 mM Glucose (Sigma Aldrich), 2 mM L-Glutamine (Gibco), 20 mM HEPES (Gibco) 500 nM SYTOX Green (Thermo Fischer) and 2.5 µM DRAQ5 (Biostatus) and seeded at a density of 5x10^5^ cells per well into µ-slide 8 well ibiTreat dishes (ibidi). Cells were stimulated with final concentrations of 100 nM PMA and 10 nM PVL. Images were collected at 2 min intervals on a Leica TCS SP8 confocal microscope equipped with a climate chamber at 37°C and with a Leica HC PL APO 20x/0.75 IMM CORR CS2 objective using glycerol immersion [45].

### Mass spectrometry (MS)

Human neutrophils from three different healthy donors were seeded in 6-well tissue culture plate to a density of 3x10^5^ cells per well (SF-RPMI without HSA) and then stimulated with 50 nM PMA, 10 nM PVL, or 15 µM nigericin for 4 h to induce NETs. As a control, neutrophils were incubated in medium for 4 h. Media was carefully removed followed by a wash with fresh media to remove unbound proteins. NETs were collected by treatment with lysis buffer (1% SDS, 50 mM HEPES pH 8, 10 mM tris-(2-carboxyethyl)phosphine (TCEP), 40 mM Chloroacetamide and protease-inhibitor cocktail) and subsequent scraping.

All samples were subjected to SP3 sample preparation [69]. Briefly, proteins were denatured, reduced and alkylated, and subsequently digested with Trypsin and Lys-C proteases. TMT 11plex (Pierce) labelling was used for peptide multiplexing and quantification. Samples were mixed, desalted using solid phase extraction (Seppak 1cc/50 mg, Waters), and fractionated using basic reversed phase fractionation on a quaternary Agilent 1290 Infinity II UPLC system equipped with a Kinetex Evo-C18 column (150 x 2.1 mm, 2.6µm, 100 Å, Phenomenex). Fractions were concatenated into 8 final samples, dried down and resuspended in 2% acetonitrile, 0.1% trifluoroacetic acid (TFA) prior MS analysis. All samples were analyzed on an Orbitrap Q Exactive HF (Thermo Scientific) that was coupled to a 3000 RSLC nano UPLC (Thermo Scientific). Samples were loaded on a pepmap trap cartridge (300 µm i.d. x 5 mm, C18, Thermo) with 2% acetonitrile, 0.1% TFA at a flow rate of 20 µL/min. Peptides were separated over a 50 cm analytical column (Picofrit, 360 µm O.D., 75 µm I.D., 10 µm tip opening, non-coated, New Objective) that was packed in-house with Poroshell 120 EC-C18, 2.7 µm (Agilent). Solvent A consists of 0.1% formic acid in water. Elution was carried out at a constant flow rate of 250 nL/min using a 180-minute method: 8-33% solvent B (0.1% formic acid in 80% acetonitrile) within 120 minutes, 33-48% solvent B within 25 minutes, 48-98% buffer B within 1 minute, followed by column washing and equilibration. The mass spectrometer was operated in data-dependent acquisition mode. The MS1 survey scan was acquired from 375-1500 m/z at a resolution of 120,000. The top 10 most abundant peptides were isolated within a 0.7 Da window and subjected to HCD fragmentation at a normalized collision energy of 32%. The AGC target was set to 2e5 charges, allowing a maximum injection time of 78 ms. Product ions were detected in the Orbitrap at a resolution of 45,000. Precursors were dynamically excluded for 45 s. Raw files were processed with Proteome Discoverer 2.3 (Thermo Scientific) using SEQUEST HT for peptide identification. Peptide-spectrum-matches (PSMs) were filtered to a 1% false discovery rate (FDR) level using Percolator employing a target/decoy approach. The protein FDR was set to 1%. Further data processing was carried out in R and Perseus (v. 1.6.2.3). Only proteins identified with at least two peptides were included in the analysis. All contaminant proteins were filtered out. A three-step normalization procedure was applied. First, the total intensity of each TMT channel was normalized to correct for mixing errors. Next, the common channel in both TMT sets was used for internal reference scaling [70] in order to correct for batch effects. Afterwards the data was normalized applying trimmed mean of M values (TMM) using the edgeR package in R [71]. Proteins were filtered on those detected in at least two of three replicate experiments. Remaining undetected (NA) values were replaced with the sample-wise minimum abundance as an estimation of the limit of detection. Differential protein abundances were calculated using the limma package in R [72]. Principle component analysis and calculation of Euclidean distance between proteome samples were performed using scaled log2(abundance) values.

### NET killing assay

Purified neutrophils were seeded at 10^6^ cells per well in a flat-bottom 96-well plate. NET formation was induced with 100 nM PVL, 100 nM PMA or 30 µM nigericin for 4 h at 37 °C. If applicable, residual phagocytosis was subsequently blocked with 10 µg/ml cytochalasin B (Abcam) for 15 min. *S. aureus* (USA300) in mid-logarithmic phase was added at a multiplicity of infection of 2 in RPMI with 10% human serum (Sigma). Bacteria were spun down for 5 min at 800 x g and incubated for 1 h at 37 °C. After incubation, NETs were treated with 2 U micrococcal nuclease (Takara Bio) for 10 min at RT. Samples were resuspended, serially diluted in DPBS and plated on trypticase soy agar plates. The plates were incubated overnight at 37 °C and colony-forming units were counted. Bacterial viability was expressed as a percentage of bacteria incubated for 1h without NETs. For DNA quantification, NETs were induced as described above and digested with 2 U micrococcal nuclease for 10 min at RT, and subsequently with 0.5 mg/ml of proteinase K (Invitrogen) for 1 h at 50°C. The samples were vortexed and the DNA concentration was measured on Nanodrop 2000 spectrophotometer (ThermoFisher Scientific).

### Neutrophil markers

1x10^6^/ml neutrophils were fixed for 15 min using 4% PFA and washed to PBS supplemented with 0.1% HSA. Cells were incubated with anti-CD63-PE, anti-CD66b-APC (Miltenyi Biotec), 5 µg/ml anti-CD45-AlexaFluor 647 (Santa Cruz Biotechnology), 10 µg/ml CD88 (S5/1)-FITC (Santa Cruz Biotechnology) and 10 µg/ml anti-human C5L2-APC (BioLegend) antibodies for 30 min in the dark, washed thereafter to PBS supplemented with 0.1% HSA and measured on a CytoFLEX (BeckMan Coulter) or MACSQuant (Miltenyi Biotec).

### Patient and control characteristics

Patient demographic and clinical data are summarized in S2 Table. Controls were matched for gender and age. All study participants provided written informed consent and were free of infections at the time of blood withdrawal. The study was approved by the local Ethics committee (EA2/003/19). Healthy control samples were collected according to the approval and guidelines of the local ethics committee (EA1/0104/06).

### Statistical analysis

Analysis of differential protein abundance was performed in limma (Linear Models for Microarray data) in R, which has been shown to outperform t-tests in detecting significant changes in protein abundance [73]. Protein-wise linear models were fit and batch-corrected using the formula (0 + condition + batch) and significant changes in abundance were tested using an empirical Bayes moderated t-statistic with Benjamini-Hochberg correction. Changes in abundance were considered significant at an absolute log2 fold change > 1, and an adjusted *p* value < 0.01. Euler diagrams of significant changes across samples were visualized with the eulerr package [74]. Significant proteins were clustered by pattern across conditions usink-means clustering with indicated number of clusters. Heatmaps were produced using pheatmap package [75].

R scripts for proteome analysis and visualization can be found in the supporting zip document.

Data are represented as mean±SD unless otherwise stated.

## MS data availability

The mass spectrometry proteomics data have been deposited to the ProteomeXchange Consortium via the PRIDE [76] partner repository with the dataset identifier PXD025702.

Reviewer account details:

**Password:** Yr3cVMPZ

## Acknowledgements

We thank Christian Frese from the Proteomics Research Platform at the Max Planck Unit for the Science of Pathogens for proteomic analysis.

We thank Alf Herzig for visualizing PVL-induced NET formation in healthy and CGD neutrophils in real time by confocal microscopy.

## Author Contributions

HJ, AZ, RK, and GM designed the study

HJ, DC, and GM performed experiments and analyzed the data HVB and RK cared for the patients

AL, LGH, RL, JTS, MSS, MS, and HVB have interpreted the data and revised the manuscript for important intellectual content

CJH analyzed the mass spectrometry data HJ, AZ, RK, and GM wrote the manuscript

## Supplemental Figures

**S1 Fig.**
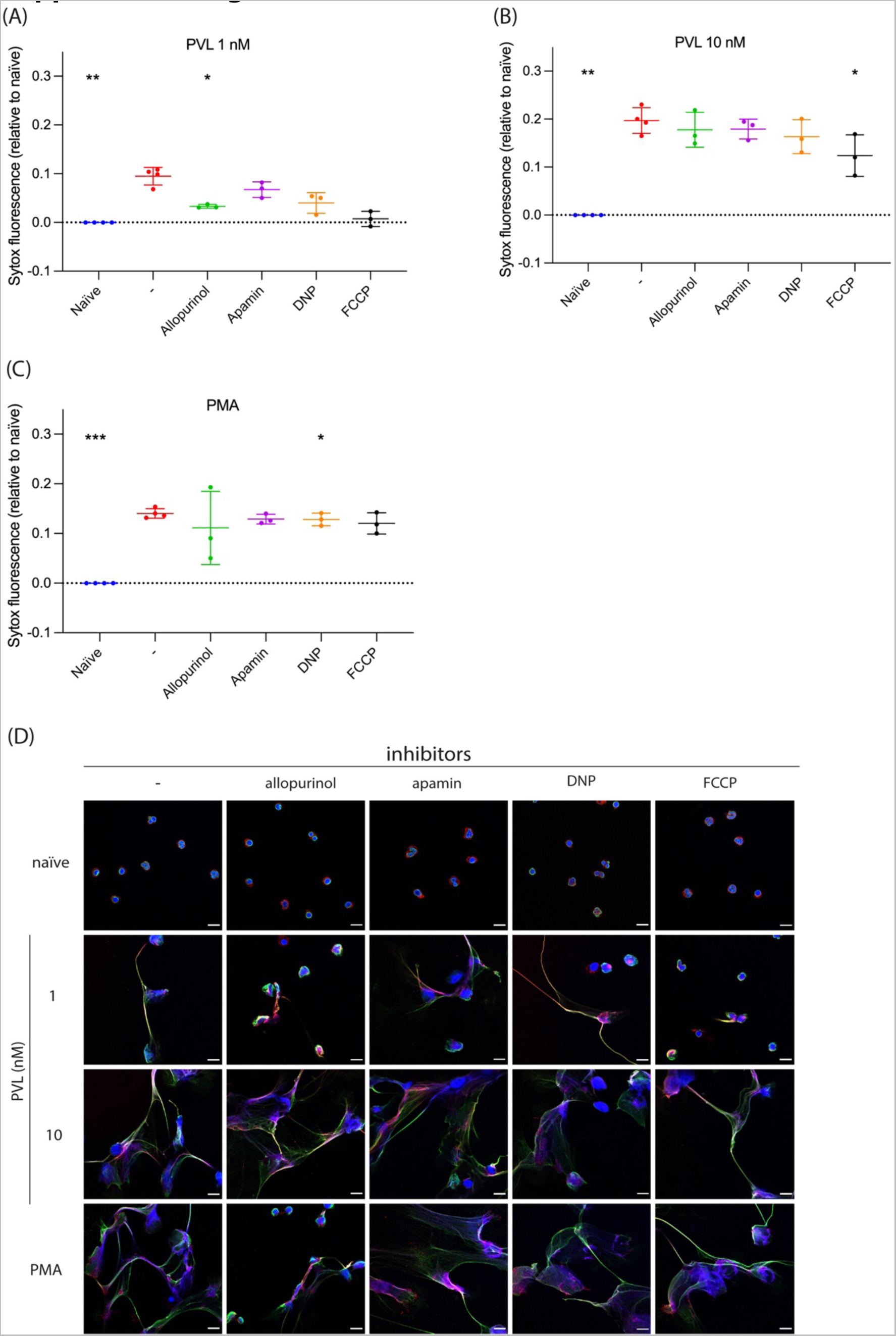
FCCP and allopurinol inhibit PVL-induced NET formation. Human primary neutrophils were treated with xanthine oxidase inhibitor allopurinol (2 mM), small conductance calcium-activated potassium channel inhibitor apamin (200 nM), and mitochondrial uncouplers DNP (750 μM) and FCCP (10 μM) for 30 min before stimulating with (A) PVL 1 nM, (B) PVL 10 nM or (C) PMA for 4 h. We quantified cell death using SYTOX Green and fluorescence relative to naïve is plotted for the respective stimuli. (D) Representative confocal microscopy of naïve neutrophils or after stimulation with PMA or PVL in the presence or absence of indicated inhibitors, and stained for DNA (blue), NE (red) and chromatin (green). Scale bar represents 10 µm. (A-C) Mean ± SD of three independent experiments. \**P*<0.05, ** *P*<0.01, *** *P*<0.001 mixed-effects analysis with Dunnett’s multiple comparison test.

**S2 Fig.**
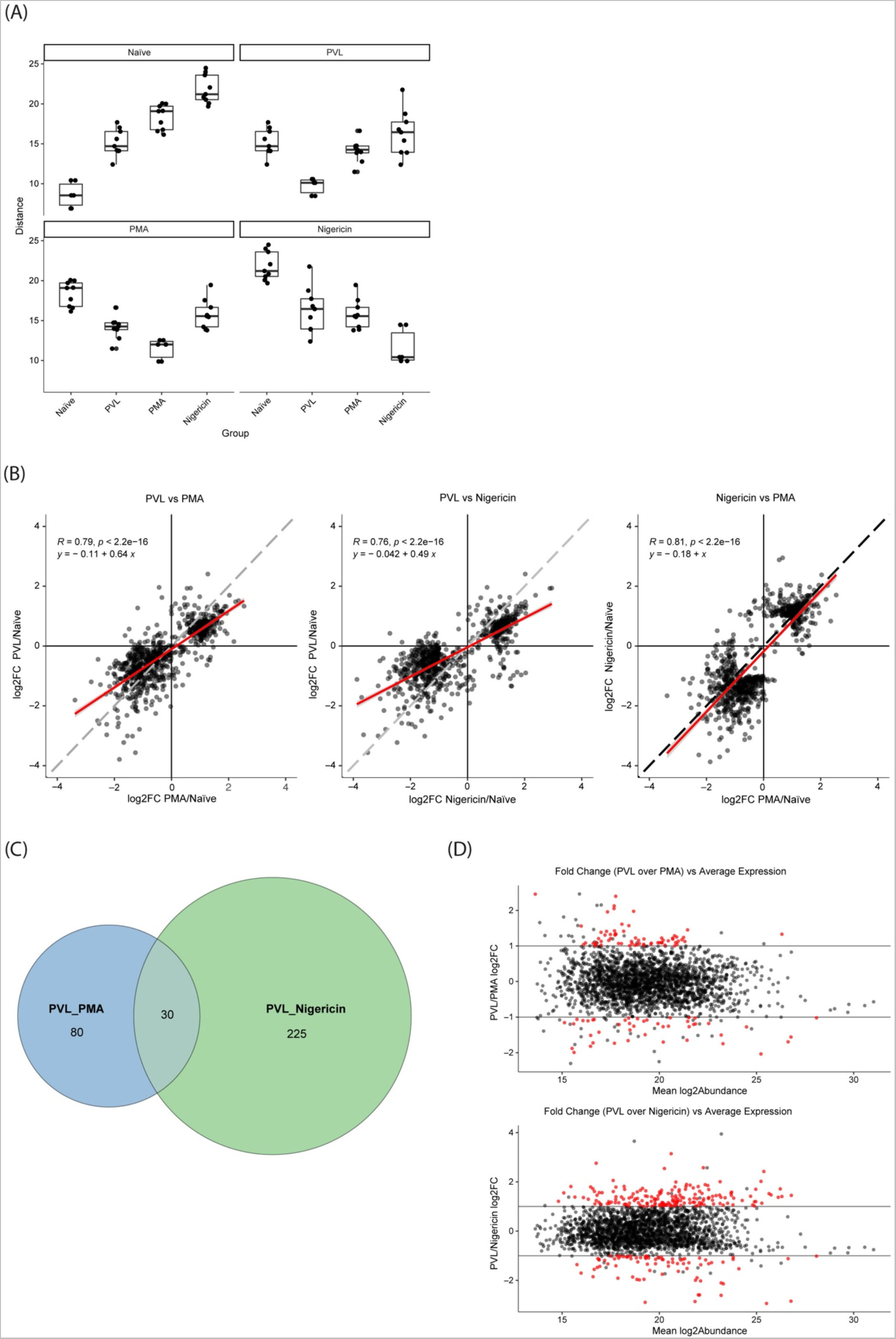
Comparison of PVL, PMA, and nigericin NET proteomes. (A) Euclidean distance between whole proteomes of naïve neutrophils and NETs induced by different stimuli. For each panel, the pairwise distance is plotted between each sample of that group and the samples of the group indicated on the x-axis. Each point represents the distance between a unique pair of samples. (B) Scatter plots of DAP fold changes compared to naïve neutrophils with linear regression for PVL vs PMA NETs, PVL vs nigericin NETs, and nigericin vs PMA NETs. DAPs plotted are proteins significantly enriched or depleted in one or more NET groups compared to naïve neutrophils (as in Fig 3B and 3C). (C) Protein log2 fold change in PVL/PMA NETs or PVL/nigericin NETs vs mean abundance across samples. Data are from pairwise comparisons between PVL and PMA or PVL and nigericin NETs. Proteome data are from n=3 donors for each condition. Significant DAPs are colored red. DAPs from each indicated comparison were considered significant with an absolute log2 fold change > 1 and adjusted *p* value < 0.01.

**S3 Fig.**
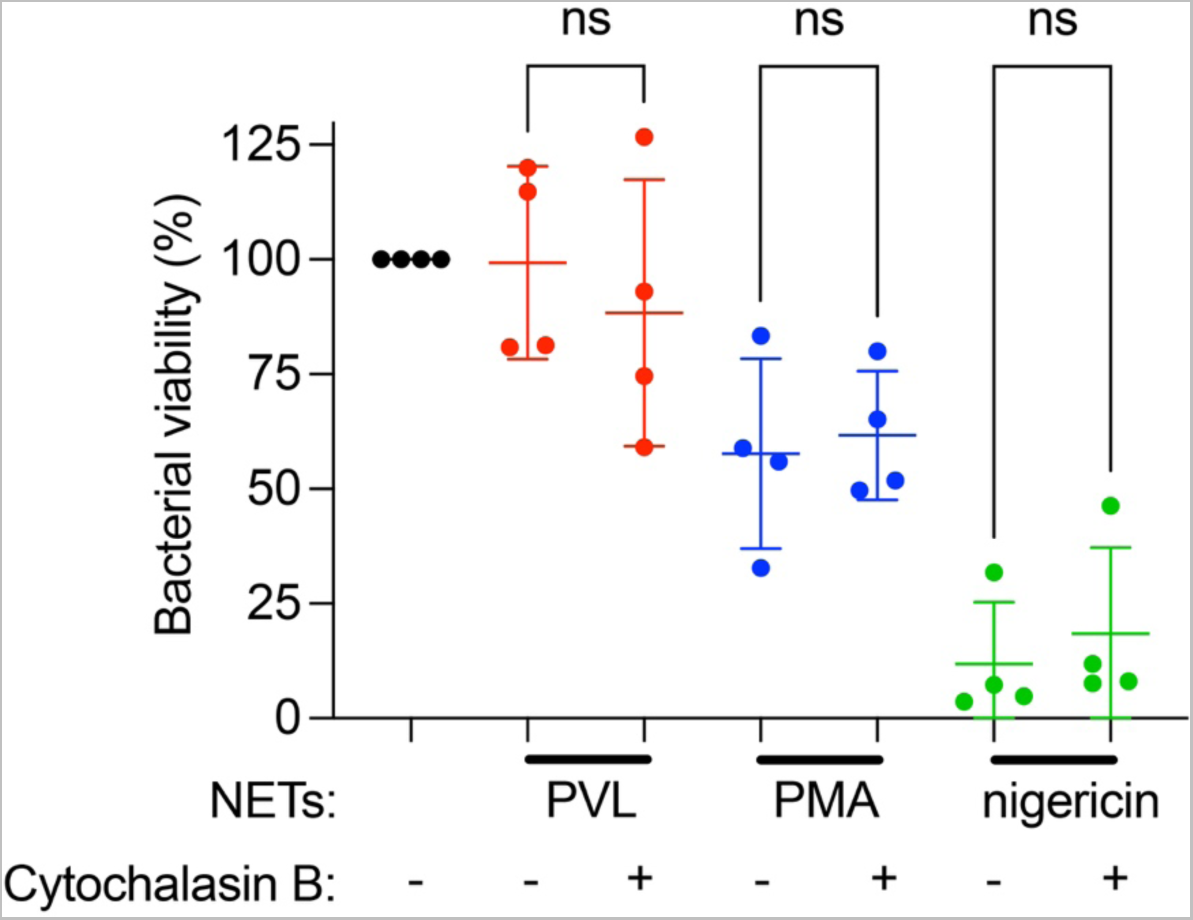
Inhibition of phagocytosis by cytochalasin B does not affect killing by NETs. NET formation was induced by PVL (100 nM), PMA (100 nM), or nigericin (30 µM) for 4 h, followed by incubation with cytochalasin B (10 µg/ml) for 15 min. MRSA was incubated with the NETs for 1 h and CFUs were quantified after overnight incubation on TSA plates. Results are expressed as a percentage of CFUs formed by MRSA incubated in the absence of NETs.

**S4 Fig.**
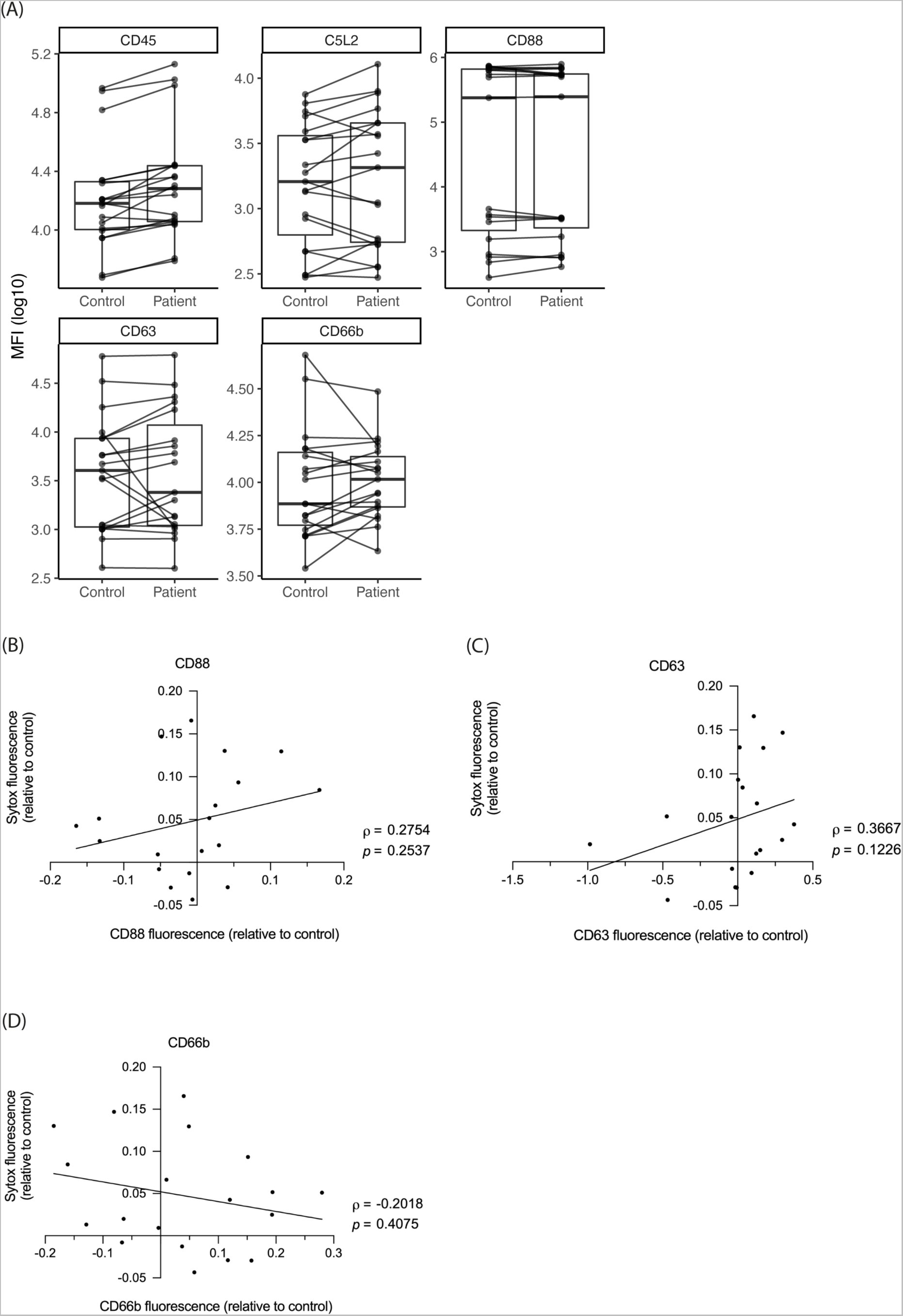
Comparison of receptor expression between patients and controls and correlation with cell death. (A) Pair-wise comparison of CD45, C5L2, CD88, CD63, and CD66b MFI between patients and controls. (B-D) CD88, CD63, and CD66b fluorescence relative to control was plotted against SYTOX Green fluorescence induced by 0.1 nM PVL relative to control. The coefficient of correlation (ρ) and probability (p) of a Spearman’s correlation test are indicated. A best-fit line indicates the trend.

## S1-8 movies

Live cell imaging of control or CGD neutrophils left untreated (S1 and S5) or stimulated with 50 nM PMA (S2 and S6) or 1 nM PVL (S3-S4 and S7-S8) over a time course of 6 h. Images were acquired every 2 min. Cells were stained with the cell permeable DNA dye DRAQ5 (magenta) and the cell impermeable DNA dye SYTOX Green (green).

## List of abbreviations

CFU: Colony forming unit
CGD: Chronic granulomatous disease
DAP: Differentially abundant proteins
DAPI: 4’6-diamidino-2-phenylindole
DHR: Dihydrorhodamine
DNP: Dinitrophenol
DPBS: Dulbeccos’s phosphate-buffered saline
DPI: Diphenyleneiodonium chloride
FCCP: Carbonyl cyanide-p-trifluoromethoxyphenylhydrazone
FDR: False discovery rate
HCD: Higher-energy collisional dissociation
HEPES: Hydroxyethyl-Piperazine-Ethane Sulfonic Acid
Hlg: γ-Hemolysin
HAS: Human serum albumin
MFI: Mean fluorescent intensities
MPO: Myeloperoxidase
MRSA: Methicillin-resistant *Staphylococcus aureus*
MS: Mass spectrometry
NE: Neutrophil elastase
NEi: Neutrophil elastase inhibitor
NETs: Neutrophil extracellular traps
PBS: Phosphate-buffered saline
PFA: Paraformaldehyde
PKC: Protein Kinase C
PMA: Phorbol 12-myristate 13-acetate
PMN: Polymorphonuclear leukocytes
PSMs: Peptide-spectrum-matches
PVL: Panton-Valentine leukocidin
PVL-SA: PVL-positive *S. aureus*
ROS: Reactive oxygen species
RT: Room temperature
S. aureus: Staphylococcus aureus
SDS: Sodium Dodecyl Sulfate
SSTI: Skin and soft tissue infections
TCEP: Tris-(2-carboxyethyl)phosphine
TFA: Trifluoroacetic acid
TMM: Trimmed mean of M values
TMT: Tandem Mass Tag

