## Supplementary Table 2 for "Panton-Valentine leukocidin-induced neutrophil extracellular traps lack antimicrobial activity and are readily induced in patients with recurrent PVL^+^-*Staphylococcus aureus* infections"

| Code | Patient | Age | Gender | Ethnicity | *S. aureus* | Symptoms | Underlying condition | Relation to other patients |
|---|---|---|---|---|---|---|---|---|
| **1942.001** | 1 | 77 | male | Caucasian | MRSA | recurrent abscesses | - | |
| **1983.003** | 2 | 36 | male | African | MRSA | recurrent abscesses | - | |
| **1957** | 3 | 62 | male | Caucasian | MSSA | severe, multifocal SSTI, necrotizing fasciitis, sepsis | obesity, diabetes | |
| **1982.002** | 4 | 37 | female | Arab | MRSA | recurrent abscesses | - | mother of patient 5 |
| **2001.001** | 5 | 14 | male | Arab | MRSA | recurrent abscesses, mastoiditis, lymphadenitis, sinus vein thrombosis | - | son of patient 4 |
| **1997.004** | 6 | 22 | male | Caucasian | MSSA | recurrent abscesses, hordeola | - | partner of patient 7 |
| **1998.020** | 7 | 22 | female | Caucasian | MSSA | recurrent abscesses, hordeola | - | partner of patient 6 |
| **1990.005** | 8 | 30 | female | Caucasian | MRSA | recurrent abscesses | - | partner of patient 9 |
| **1989.006** | 9 | 30 | male | Caucasian | MRSA | recurrent abscesses | - | partner of patient 8 |
| **2006.007** | 10 | 13 | male | Arab | MSSA | recurrent abscesses, hordeolum | - | |
| **1975.014** | 11 | 45 | male | Caucasian | MSSA | recurrent severe SSTI | - | father of patient 12<br>son of patient 13 and 14 |
| **2009.016** | 12 | 14 | female | Caucasian | MSSA | recurrent abscesses | - | daughter of patient 11<br>granddaughter of patients 13 and 14 |
| **1952.018** | 13 | 68 | male | Caucasian | MSSA | recurrent abscesses | - | father of patient 11, grandfather of patient 12 |
| **1953.19** | 14 | 67 | female | Caucasian | MSSA | recurrent abscesses | - | mother of patient 11, grandmother of patient 12 |
| **1986.004** | 15 | 34 | male | Caucasian | MSSA | recurrent abscesses | - | partner of patient 16 |
| **1984.005** | 16 | 36 | female | Caucasian | MSSA | one abscess | - | partner of patient 15 |
| **1963.006** | 17 | 57 | male | Caucasian | MRSA | recurrent abscesses | - | - |

| | | | | | | | | |
|---|---|---|---|---|---|---|---|---|
| **1975.021** | 18 | 45 | female | Caucasian | MSSA | recurrent abscesses | - | - |
| **2001.001** | 19 | 19 | male | Caucasian | MSSA | necrotizing pneumonia, septic thrombembolism | Influenza | - |
